# Fast4DReg: Fast registration of 4D microscopy datasets

**DOI:** 10.1101/2022.08.22.504744

**Authors:** Joanna W Pylvänäinen, Romain F Laine, Bruno M.S. Saraiva, Sujan Ghimire, Gautier Follain, Ricardo Henriques, Guillaume Jacquemet

## Abstract

Unwanted sample drift is a common issue that plagues microscopy experiments, preventing accurate temporal quantification of biological processes. While multiple methods and tools exist to correct images post-acquisition, performing drift correction of large 3D videos using open-source solutions remains challenging and time-consuming. Here we present a new tool developed for ImageJ/Fiji called Fast4DReg that can quickly correct axial and lateral drift in 3D video microscopy datasets. Fast4DReg works by creating intensity projections along multiple axes and estimating the drift between frames using 2D cross-correlations. Using synthetic and acquired datasets, we demonstrate that Fast4DReg performs better than other state-of-the-art open-source drift correction tools and significantly outperforms them in speed (5x to 60x). We also demonstrate that Fast4DReg can be used to register misaligned channels in 3D using either calibration slides or misaligned images directly. Altogether Fast4DReg provides a quick and easy-to-use method to correct 3D imaging data before further visualization and analysis. Fast4DReg is available on [GitHub](https://github.com/guijacquemet/Fast4DReg).

## Introduction

Live imaging is essential in biomedical research, enabling scientists to follow biological processes over time. Despite being heavily used, performing live imaging experiments using fluorescence microscopy remains technically challenging. The experimenter must carefully balance illumination power and acquisition speed while maintaining specimen health. In addition, imaging is often prone to drift. Drift can occur due to the microscope instability caused, for example, by temperature changes leading to thermal expansion of the mechanics or by the movement of the sample itself.

Multiple software and hardware solutions have been developed to minimize drifting during acquisition. For instance, axial drifting can be limited using an infrared light that is reflected on the glass-sample interface and captured by a detector (e.g., Leica’s Adaptive Focus Control or Nikon’s Perfect Focus System). Lateral drift due to sample movement can also be compensated by tracking algorithms that follow the sample over time and move the microscope stage accordingly (Fox et al., 2022; von Wangenheim et al., 2017). Yet drifting is rarely entirely eliminated at the acquisition stage, especially when acquiring multiple positions for an extended period. Therefore, it is often necessary to perform drift correction (via image registration) as a post-processing step before image visualization and quantification. Beyond live imaging, drift correction/channel registration is a crucial processing step for multiple image analysis pipelines, including colocalization analysis or the reconstruction of super-resolution microscopy images.

Most drift correction/registration algorithms work sequentially by comparing a reference image to a moving image and estimating the movement between these two images to correct it. Multiple open-source tools capable of correcting 4D datasets already exist. Popular tools include, for instance, ITK (McCormick et al., 2014), elastix (Klein et al., 2010), Multiview Reconstruction (Preibisch et al., 2010, 2014), Fijiyama (Fernandez Moisy, 2021), or Correct 3D drift (Parslow et al., 2014). However, except for Multiview Reconstruction and Correct 3D drift, these tools are geared toward correcting medical imaging datasets and can be unpractical to use to correct large 3D movies. Multiview Reconstruction, which was designed to register large lightsheet fluorescence microscopy datasets, uses beads and/or segmented structures in the imaging volume to perform the registration, which are not always available (Preibisch et al., 2010, 2014). While Correct 3D drift can often successfully register our datasets, we felt limited by its speed and available features.

Here, prompted by a need to correct our 3D videos more easily and more efficiently, we developed Fast4DReg, a fast 2D and 3D video drift correction tool. Using multiple datasets, we show that Fast4DReg can perform better than two other state-of-the-art 3D video drift correction tools available in Fiji, namely Correct 3D drift (Parslow et al., 2014) and Fijiyama (Fernandez and Moisy, 2021). In addition, we show that Fast4DReg can register misaligned channels in 3D using either calibration slides or misaligned images directly. Fast4DReg is fast and has a simple graphical interface. These features make Fast4DReg a versatile and easy-to-use open-source 2D/3D drift correction tool.

## Implementation, operation, and test datasets

### Pipeline

Fast4DReg breaks the drift correction task into two steps: First, estimation of drift followed by applying the drift correction. To estimate the drift of a 3D video in x,y, and z, Fast4DReg sequentially estimates the lateral drift, corrects the lateral drift, and then estimates and corrects the axial drift (Figure 1). Lateral and axial drift corrections can also be performed independently, which can be particularly useful when only axial drift needs to be corrected. As an output of the drift estimation step, Fast4DReg creates a new folder containing the corrected images, the drift plots, a drift table, and a settings file containing the selected parameters. Notably, the drift table can then be applied to correct other images using the same parameters (i.e., to correct another channel). Indeed, when correcting multichannel 3D videos, the user needs to choose one channel to use to estimate the drift. The other channel(s) can then be corrected using the same drift table (Figure 1).

**Fig. 1.**
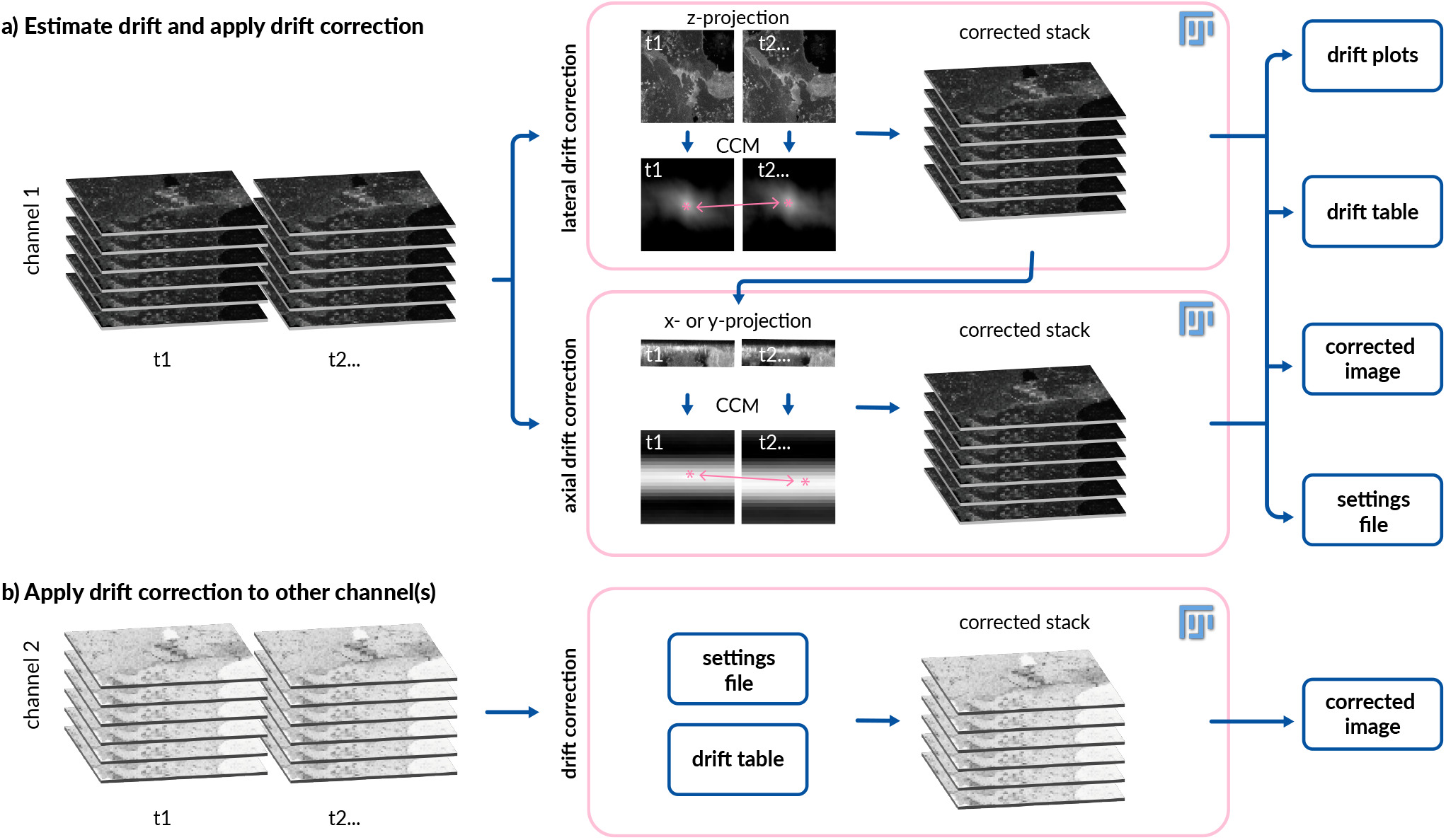
Drift correction of 3D videos using Fast4DReg. Scheme highlighting the inner working of Fast4DReg. **a**) Fast4DReg sequentially estimates the lateral drift, corrects the lateral drift, and then estimates and corrects the axial drift. Fast4DReg creates intensity projections along multiple axes and estimates the drift between the reference and moving frames by calculating their cross-correlation matrix (CCM). The location of the peak intensity in the CCM (pink asterisk) defines the linear shift between the two images (as highlighted by the pink arrow). Fast4DReg outputs the corrected images, the drift plots, a drift table, and a settings file containing all selected parameters and paths to the drift table. **b**) The settings file inducing the used parameters and path to the drift table can then be applied to correct other datasets (i.e., another channel) directly.

To estimate the lateral or axial drift of a 3D video, Fast4DReg creates z or y intensity projections for each time point to create a 2D video. Fast4DReg then estimates the linear drift between the reference and moving frames by calculating their cross-correlation matrix (CCM). The location of the peak intensity in the CCM defines the linear shift between the two images. Sub-pixel accuracy is accomplished by up-scaling the CCM via bicubic spline interpolation [as demonstrated by (Laine et al., 2019)]. Depending on their data, users can choose the first frame (best for fixed data) or the movie’s previous frame (best for live imaging data) as the reference frame.

Fast4DReg can also be used to register channels from misaligned 3D stacks. In this case, Fast4DReg simply converts the channels into time frames before applying the Fast4DReg drift correction pipeline described above. As a note of caution, cross-correlation only works well to register channels where similar structures/cells are labeled.

Importantly Fast4DReg can also register 2D video and 2D multichannel images either one at a time or in batch.

#### Operation

Fast4DReg can run on any computer where Fiji (Schindelin et al., 2012) and the Bio-Formats (Linkert et al., 2010) plugin are installed. Fast4DReg has a RAM-saving mode that allows the registration of large datasets using a computer with limited resources at the cost of a slightly longer processing time.

Fast4DReg expects as input one or multiple single channel 2D or 3D video and outputs corrected files, drift tables, drift plots, and a settings file that can be applied to other channels as needed. Thanks to Bio-Formats (Linkert et al., 2010), Fast4DReg can handle various image formats as input. Fast4DReg can be tested using our test datasets available on [Zenodo](https://zenodo.org/record/6534570). Installation procedure and step-by-step instructions are available on the Fast4DReg [GitHub](https://github.com/guijacquemet/Fast4DReg) page.

#### Test datasets

To assess the drift correction ability of Fast4DReg, we used four types of datasets: (1) Synthetic drift -datasets, (2) a HUVEC monolayer -dataset, and (3) a registration slide -dataset, and (4) a Filopodia-dataset. These datasets and the code generated are available on a dedicated [Zenodo](https://zenodo.org/record/6534570) archive.

#### Dataset 1: Datasets with synthetic drift

Synthetic drift 3D video datasets were created by duplicating 25 times a 3D stack image (416×416 px, 69 z-slices, 25-time points, 16 bit) and artificially adding a certain amount of x-, y- and z drift between each frame (Figure 2a). The amount of drift added corresponds to drift typically observed in our live-cell imaging experiments. Using this method, three videos were created: One with no drift (ground truth video), one with only a small amount of drift, and one with a large drift (across the field of view). The videos were created using the following functions:

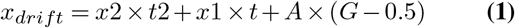

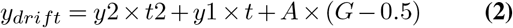

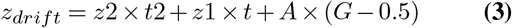

**Fig. 2.**
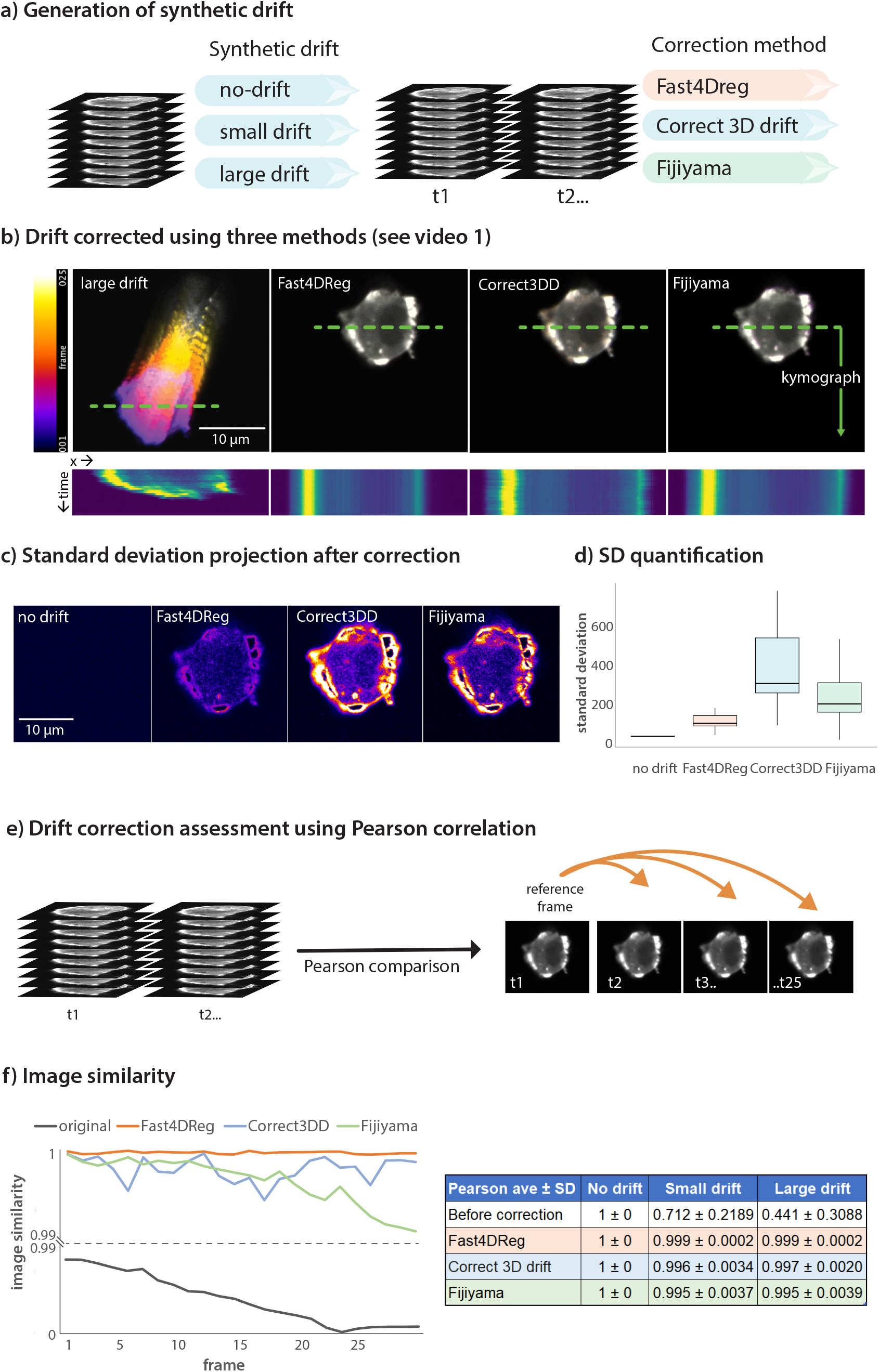
Figure 2: Fast4DReg outperforms Correct 3D drift and Fijiyama on a synthetic 3D + t dataset. **a**) Three synthetic 3D video datasets were created, one with no drift, one with a small amount of drift, and another with a large amount of drift. The small and large drift datasets were then corrected using Fast4DReg, Correct 3D drift, and Fijiyama. Of note, the settings giving the best performance were used for each tool (see methods for details). **b**) The drift correction performance of the three algorithms was first visually assessed using a temporal color projection of the middle slice of the stack and a kymograph (along a white dashed line). **c**) Standard deviation projection of the middle slice of the stack. This projection takes the standard deviation of the pixel intensities through the stack. Positions with large differences in the pixel intensities through the stack appear brighter in this projection. Therefore a black image highlight no variation between the frames over time (perfect drift correction), while signals highlight slight errors in the drift correction. **d**) For each z-slice, the standard deviation projection over-time was generated and quantified using Fiji, and the results are shown as boxplots created by PlotsOfData (Postma and Goedhart, 2019). No drift shows a high baseline value as specified noise was added during background homogenization. **e**-**f**) Pearson’s correlation coefficient was calculated between the first and subsequent frames. A value of 1 indicates perfect drift correction. For all panels, the scale bar = 10 μm.

Where A represents the noise amplitude and G is a Gaussian “normally” distributed pseudorandom number with mean 0.0 and standard deviation 1.0.

##### No drift dataset

x1 = 0, x2 = 0, y1 = 0, y2 = 0, z1 = 0, z2 = 0, noise amplitude A = 0.0. Small drift dataset: x1 = 0.1, x2 = 0.05, y1 = 0.5, y2 = −0.05, z1 = −0.3, z2 = 0, noise amplitude A = 0.1. Large drift dataset: x1 = 0.2, x2 = 0.1, y1 = 1.5, y2 = −0.03, z1 = −0.6, z2 = 0, noise amplitude A = 0.1.

After the drifting was simulated, the image background was made homogeneous via pixel intensity subtraction and by adding specified noise using Fiji (subtract = 800, add = 800, the addition of specified noise = 100). The x-y-and z drift in these synthetic datasets was corrected (considering the whole image, not a selected ROI) using Fast4DReg, Correct 3D drift, and Fijiyama. For each software, the parameters providing the best possible drift correction were chosen and were as follows: Fast4DReg: xy-projection type = max intensity, xy time averaging = 1, xy max expected drift = disabled, xy-reference previous frame (better for live), z-projection type = max intensity, z-reslice mode = Top, z time averaging = 1, z max expected drift =disabled, z-reference previous frame (better for live), expand stack to fit = disabled, Save RAM = disabled

##### Correct 3D drift

Channel for registration = 1, Multi time scale computing = enabled, Sub pixel drift correction = enabled, Edge enhancement = enabled, all pixels and z-planes considered, max shift in x-y and z-directions = 200 px, use of virtual stack = disabled, only compute vectors = disabled).

##### Fijiyama

Max subsampling factor = 4, Min subsampling factor = 2, Higher accuracy = yes, Block half-size x= 5, y= 5, z= 3, Block neighborhood x = 2, y = 2, z = 2, Striding along x = 13, y=13,z = 3, number of iterations = 6, percentage of blocks selected by score = 95%, Percentage kept in Least-trimmed square = 80%.

After correcting the drift in the synthetic datasets, the images were first cropped to be the same size (352×275px, 69 z-slices, 25 frames) using Fiji. The drift correction performance was then quantified by measuring the Pearson’s correlation coefficient between frames (reference frame = first) of a selected z-slice (z-slice = 51) using a custom-made Jupyter notebook. This z-slice was selected as it is in the middle of the stack.

#### Dataset 2: the HUVEC monolayer -dataset

The HUVEC monolayer -dataset consists of a 3D timelapse of human umbilical vein endothelial cells (HUVECs) labeled with sirActin (Spirochrome). The video was acquired using a laser scanning confocal microscope LSM880 (Zeiss) equipped with an Airyscan detector (Carl Zeiss) and a 63x oil (NA 1.4) objective. The microscope was controlled using Zen Black (2.3), and the Airyscan was used in standard super-resolution mode. This dataset has 200 frames (488 × 488 px) and 24 z-slices (2,3 GB). This dataset suffers from significant drift in all x-, y- and z-directions.

The xy- and z-drift in this dataset was corrected (considering the whole image, not a selected ROI) using Fast4DReg, Correct 3D drift, and Fijiyama. For each software, the parameters providing the best possible drift correction were chosen and were as follows:

##### Fast4DReg

xy-projection type = max intensity, xy time averaging = 1, xy max expected drift = disabled, xy-reference = previous frame (better for live), z-projection type = max intensity, z-reslice mode = Top, z-time averaging = 1, z-max expected drift =disabled, z-reference = previous frame (better for live), expand stack to fit = disabled, Save RAM = disabled.

##### Correct 3D drift

Channel for registration = 1, Multi time scale computing = enabled, Sub pixel drift correction = enabled, Edge enhancement = enabled, all pixels and z-planes considered, max shift in x-y and z-directions = 200 px, use of virtual stack = disabled, only compute vectors = disabled).

After drift correcting this dataset, the correction performance was quantified by measuring the Pearson’s correlation coefficient between adjacent frames (reference frame = previous) of a selected z-slice (z-slice = 8) using a custom-made Jupyter notebook. Two computers were used to compare the execution times of all compared methods; Computer 1 (C1) (operating system: Windows, processor: AMD Ryzen 7 5800X 8-Core Graphics card: GeForce GTX™ 3080, RAM: 32 GB, Fiji version 1.53q) and Computer 2 (C2) (operating system: Mac, Processor: M1 chip (8-core CPU, 8-core GPU), RAM: 16 GB, Fiji version 1.53q).

The settings used to process this dataset with Fijiyama were as follows: Max subsampling factor = 4, Min subsampling factor = 2, Higher accuracy = yes, Block half-size x= 5, y= 5, z= 3, Block neighborhood x = 2, y = 2, z = 2, Striding along x = 13, y=13,z = 3, number of iterations = 6, percentage of blocks selected by score = 95%, Percentage kept in Least-trimmed square = 80%.

#### Dataset 3: The Registration slide -dataset

The Registration slide -dataset was created by imaging a channel calibration slide (Argolight ™ HM) with four channels using a widefield microscope (1024×1024 px, 25 z-slices, 4 channels). This dataset was acquired using a DeltaVision OMX v4 (GE Healthcare Life Sciences) microscope fitted with a 60x Plan-Apochromat objective lens, 1.42 NA (immersion oil RI of 1.516) used in widefield illumination mode. Emitted light was collected on a front-illuminated pco.edge sCMOS (pixel size 6.5 mm, readout speed 95 MHz; PCO AG) controlled by SoftWorx.

The x-y- and z-drift in this dataset was corrected (considering the whole image, not a selected ROI) using Fast4DReg using the following parameters: xy-projection type = max intensity, xy-time averaging = 1, xy-max expected drift = disabled, xy-reference = first frame (default, better for fixed), z-projection type = max intensity, z-reslice mode = Top, z time averaging = 1, z-max expected drift =disabled, z-reference = first frame (default, better for fixed), expand stack to fit = disabled, Save RAM = disabled.

#### Dataset 4: Filopodia-dataset

The Filopodia-dataset (1024×1024 px, 17 z-slices, 3 channels) consists of a 3D structured illumination microscopy (SIM) image of a U2OS cell expressing a GFP-tagged Lamellipodin fragment, MYO10-mScarlet, and labeled to visualize its actin cytoskeleton using sirActin (Spirochrome) (Miihkinen et al., 2022). This dataset was acquired using a DeltaVision OMX v4 (GE Healthcare Life Sciences) microscope fitted with a 60x Plan-Apochromat objective lens, 1.42 NA (immersion oil RI of 1.516) used in SIM illumination mode (five phases x three rotations). Emitted light was collected on a front-illuminated pco.edge sCMOS (pixel size 6.5 mm, readout speed 95 MHz; PCO AG) controlled by SoftWorx.

The x-y- and z-drift in this dataset was corrected (considering the whole image, not a selected ROI) using Fast4DReg using the following parameters: xy-projection type = max intensity, xy-time averaging = 1, xy-max expected drift = disabled, xy-reference = previous frame (better for live), z-projection type = max intensity, z-reslice mode = Top, z-time averaging = 1, z max expected drift =disabled, z-reference = first frame (default, better for fixed), expand stack to fit = disabled, Save RAM = disabled.

## Use cases

### Fast4DReg outperforms Correct 3D drift or Fijiyama on our synthetic dataset

To assess the capabilities of Fast4DReg to correct 3D videos, we compared Fast4DReg results to two other widely used state-of-the-art drift correction methods available in Fiji (Schindelin et al., 2012), namely Correct 3D drift (Parslow et al., 2014) and Fijiyama (Fernandez and Moisy, 2021). For this purpose, three synthetic videos with known amounts of drift were created: one with no drift, one displaying a small amount of drift, and another with a larger amount of drift (Figure 2a and the Test datasets section for details). As these videos were generated by duplicating an acquired single 3D stack and adding artificial drift, a perfect drift correction will generate identical time frames.

Visually, all three tools successfully corrected the artificially drifting 3D videos regardless of the amount of drift (Figure 2b, Video 1). To carefully quantify the performance of these three software, we selected a z-slice and (1) plotted the standard deviation projection of the corrected stack (Figure 2c and 2d), and (2) calculated the Pearson’s correlation coefficient between the first and each subsequent frames (Figure 2e and 2f). Both assessment methods indicate that Fast4DReg performs slightly better than Correct 3D drift or Fijiyama on our synthetic dataset (Video 2, Figure 2c-2f). Importantly, these results demonstrate that 2D intensity projection followed by 2D cross-correlation is a viable method to correct drifting 3D videos.

#### Fast4DReg is fast and efficiently corrects drift from acquired 3D videos

Next, we assessed the suitability of Fast4DReg to correct drifts in acquired 3D biological images. We used a long 3D video of a HUVEC monolayer labeled with sir-Actin that suffered from significant xyz-drifting during the imaging. We also registered this dataset with Correct 3D drift and Fijiyama. While both Fast4DReg and Correct 3D drift produced good results when assessed visually (Video 3), we failed to generate meaningful results with Fijiyama as the processing made the video drift even more than the raw data (data not shown).

To estimate the correction efficiency of Fast4DReg and Correct 3D drift on this dataset, we first searched for a structure that should remain immobile across multiple time points in the movie and chose a large stress fiber. We then color-coded three consecutive frames (one color per frame) and observed the overlaps of this stable structure between frames using line profiles (Figure 3a). In the uncorrected movie, the stress fiber does not overlap in these three frames, clearly indicating drift. In the movies corrected by Fast4DReg and Correct 3D drift, the stress fiber overlap between frames improves, showing that the drift correction works in both cases. Interestingly the drift correction provided by Fast4DReg appears superior here as the stress fiber overlap between the three frames is greater despite the correction being performed globally (and not locally to that specific structure) (Figure 3a).

**Fig. 3.**
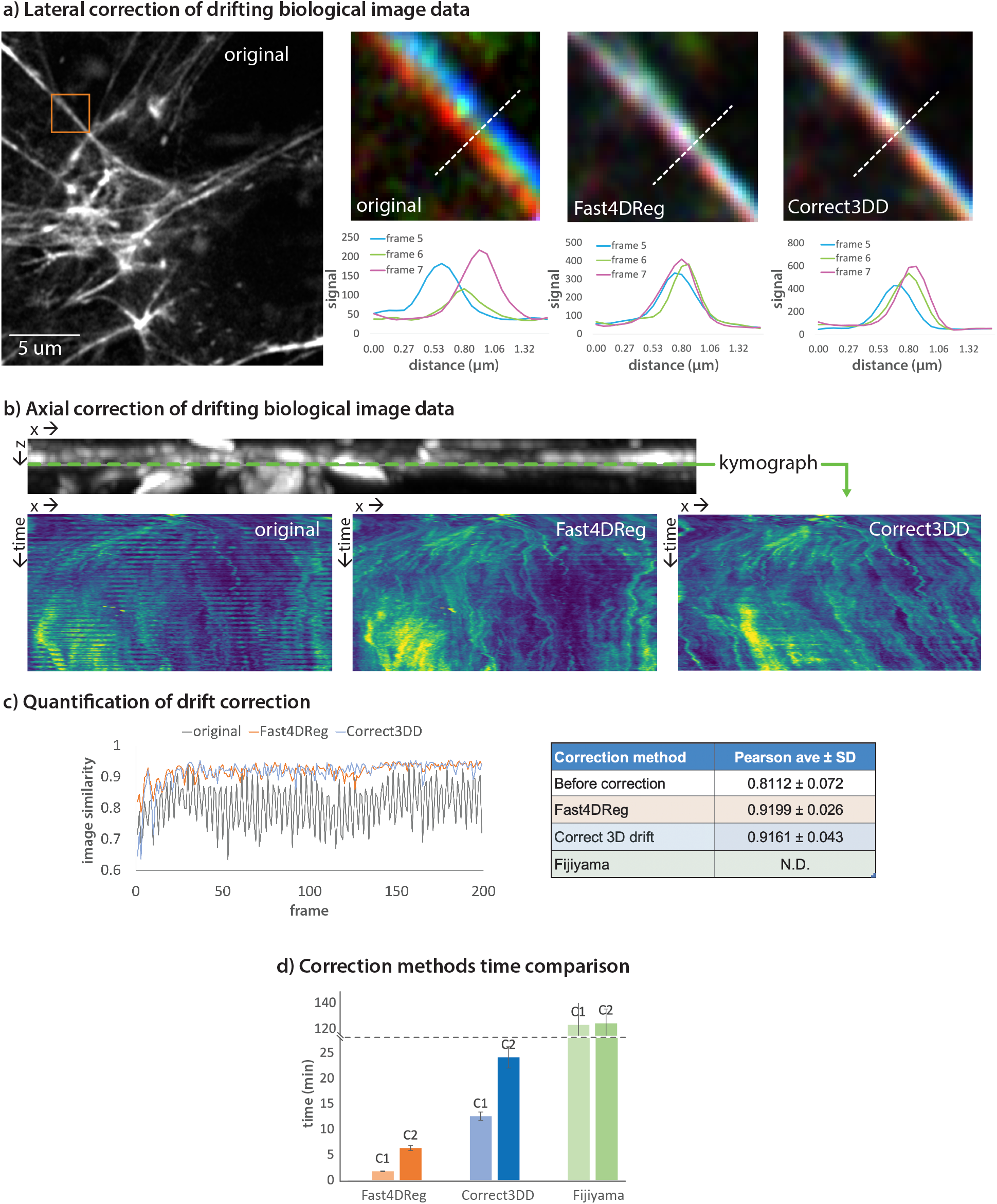
Fast4DReg can rapidly correct axial and lateral drift in 3D videos. A 3D video of HUVECs cells labeled with sirActin displaying large xyz-drift was corrected with Fast4DReg and Correct 3D drift. **a**) A region of interest containing a stress fiber that should remain immobile across multiple time points was chosen. Three consecutive frames were pseudo-colored red, green, and blue and merged. White indicates structural overlaps between the three frames. Line profiles to further study the overlap between frames were drawn as shown. **b**) Kymograph of a selected line in y-projection was created to visualize the drift in z over time. **c**) Pearson’s correlation coefficient was calculated between each consecutive frame pair. **d**) Two computers, computer 1 (high-performance desktop) (C1) and computer 2 (laptop) (C2) were used to measure the speed of the correction methods. Shown values are the average times of three measurements (C1, operating system: Windows, processor: AMD Ryzen 7 5800X 8-Core Graphics card: GeForce GTX™ 3080, RAM: 32 GB, Fiji version 1.53q; C2, operating system: Mac, Processor: M1 chip 8-core CPU, 8-core GPU, RAM: 16 GB, Fiji version 1.53q).

To visualize the axial drift correction efficiency of Fast4DReg and Correct 3D drift on this dataset, we generated kymographs from the y-projections (Figure 3b). In the original data, the kymograph shows a clear band pattern due to the microscope stage jumping cyclically. This banding pattern is improved in the movies corrected by Fast4DReg and Correct 3D. Still, it has not entirely disappeared, indicating that while both registration methods work well on this dataset, the correction is not perfect (Figure 3b and Video 3). This is perhaps because part of the data goes out of the imaging volume several times. Of note, in this case, Correct 3D drift processing leads to the monolayer slowly sinking over time.

To obtain a more quantitative estimate of the performance of Fast4DReg and Correct 3D drift on the HUVEC dataset, we measured Pearson’s correlation coefficients between each adjacent frame pair for a selected z-plane in the corrected videos. Indeed, we consider that efficient drift correction will make successive frames more similar to one another despite the inevitable biological changes. Using this metric, we find that Fast4DReg performed equally or slightly better than Correct 3D drift on this dataset (Figure 3c).

Our primary motivation behind developing Fast4DReg was to create a 3D registration pipeline that is easy to use, flexible, and very fast. Therefore we next assessed the computing time required by Fast4DReg, Correct 3D-Drift, and Fijiyama to process the HUVEC dataset using two different computers. We found that Fast4DReg (1 min 48s to 6 min 24s) is 4-7 times faster than Correct 3D drift (12 min 30 to 24 min) and 20 to 70 times faster than Fijiyama (1 h to 2 h) when correcting the HUVEC dataset (Figure 3d). These differences are significant as datasets’ registration often requires tweaking hyperparameters to obtain the best possible results. Fast4DReg speed is likely due to two factors: (1) using 2D projections greatly simplifies the computations required, and (2) using CPU multithreading further accelerates the 2D cross-correlation process. Overall, Fast4DReg outperforms Correct 3D-Drift when correcting the HUVEC dataset and is much faster.

#### Fast4DReg can also register misaligned 3D channel stacks

Next, we tested if Fast4DReg could be used to align 3D multi-channel images instead of 3D videos. In this case, Fast4DReg uses the same pipeline as described for time series but first converts the channels into time frames.

To test this approach, we first registered, using Fast4DReg, a four-channel 3D image of a calibration slide imaged using a widefield microscope. In this dataset, the raw images display significant xy and z misalignment due to chromatic aberrations and the fact that the channels were acquired using different cameras (Figure 4a and 4b). Using line intensity profiles, we found that Fast4DReg can efficiently register this dataset laterally and axially (Figure 4a and 4b). As Fast4DReg can apply saved drift correction tables, we envision that the approach described here can be used to register any microscopy images when the shift is first estimated using a registration calibration slide or multi-color bead images.

**Fig. 4.**
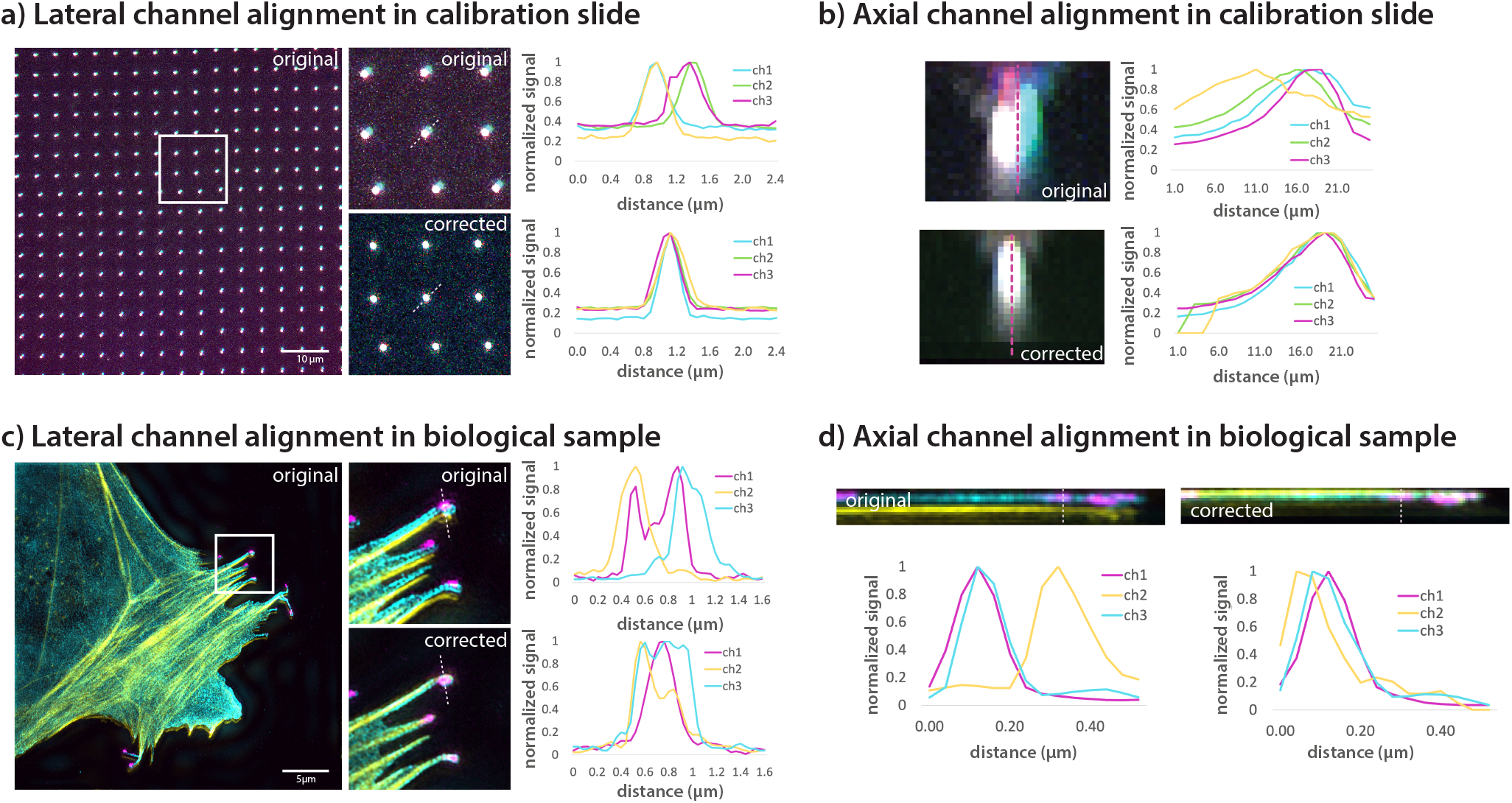
Fast4DReg can align 3D multi-channel images. **a-b**) 4-channel 3D registration slide image acquired using a widefield microscope was aligned using Fast4DReg. Merged images and line intensity profiles are displayed to highlight the level of overlap between the four channels. Scale bar = 10 μm. **a**) A single z-plane is displayed to illustrate the lateral misalignment corrected by Fast4DReg. **b**) A y-projection of one of the calibration slide spots is displayed to illustrate the axial misalignment corrected by Fast4DReg. **c-d**) A SIM image of a U2OS cell expressing a GFP-tagged Lamellipodin fragment (cyan), MYO10-mScarlet (magenta), and labeled to visualize its actin cytoskeleton (yellow) was aligned using Fast4DReg. **c**) A single z-plane is displayed to illustrate the lateral misalignment, evident in filopodia, corrected by Fast4DReg. **d**) A y-projection of one filopodium illustrates the axial misalignment corrected by Fast4DReg. Scale bar = 5 μm.

However, calibration slides or bead images are not always available, so we next tested the ability of Fast4DReg to correct misaligned channels directly. Here we chose three-channel 3D images acquired using structured illumination microscopy (Figure 4c and 4d). We chose images of cells displaying filopodia as these actin structures are very thin and narrow, rendering them almost two-dimensional. As with the registration slide image, Fast4DReg performs well when registering this dataset (Figure 4c and 4d). This approach will not work with all images, as the registration method used here will work only if sufficient structural overlap exists between the channels. Regardless, we envision that the approach described here can be beneficial in correcting channel misalignment when no calibration data is available.

### Discussion

Sample drifting is a significant challenge in live microscopy, and implementing post-processing drift correction pipelines is not always fast nor straightforward. Here we developed Fast4DReg, an ImageJ-based tool that can quickly correct axial and lateral drift in 2D and 3D videos. We show that Fast4DReg can outperform two open-source 3D drift correction tools on our test datasets. A significant advantage of Fast4DReg is that it can correct 3D videos in a fraction of the time compared to other tested tools and comes with an easy-to-use graphical user interface. Additionally, Fast4DReg is flexible and can be used for aligning multichannel 3D images.

Despite its performance, Fast4DReg has several limitations. (1) Fast4DReg can only perform translations when correcting a dataset. Rotation, scaling, or shearing transformations are not supported, although these should not be required to correct most time-course video or multichannel microscopy datasets. (2) The channel alignment is limited to images with structural conservation between channels or requires calibration slides or beads images to compute the shift maps.

With Fast4DReg, we demonstrate that using intensity projections followed by 2D cross-correlation is a quick and efficient way to register various multidimensional data types, including 3D videos and 3D multichannel datasets. In the future, it would be interesting to assess the suitability of using 3D cross-correlation directly to register similar images. But using 3D cross-correlation will likely impede processing times.

To promote adoption by the community, Fast4DReg is available through GitHub/Zenodo, where the pipeline, test datasets, and detailed step-by-step instructions are provided. With Fast4DReg, we hope to make the process of multidimensional data registration more straightforward and faster and, therefore, more accessible to the community.

## Data Availability

All datasets used in this study are available on [Zenodo](https://zenodo.org/record/6534570).

## Software Availability

Fast4DReg, the generator of synthetic drift, and the notebook used to make the Image similarity measurements (all under MIT licenses) are available on [GitHub](https://github.com/guijacquemet/Fast4DReg), and their source code is archived on [Zenodo](https://zenodo.org/record/6534570).

## Acknowledgements

The Cell Imaging and Cytometry Core facility (Turku Bioscience, University of Turku, Åbo Akademi University, and Biocenter Finland) and Turku Bioimaging are acknowledged for services, instrumentation, and expertise. Junel Solis and Anting Li are acknowledged for testing Fast4DReg and providing feedback. This study was supported by the Academy of Finland (G.J., 338537), the Cancer Society of Finland (G.J.), and the Solutions for Health strategic funding to Åbo Akademi University (G.J.). J.W. P. was supported by Health Campus Turku 2.0 funded by the Academy of Finland. R. F. L. was supported by an MRC Skills development fellowship (MR/T027924/1). G.F. was supported by an Academy of Finland postdoctoral fellowship (332402). R.H. is supported by Gulbenkian Foundation and received funding from the European Research Council (ERC) under the European Union’s Horizon 2020 research and innovation programme (grant agreement No. 101001332), the European Molecular Biology Organization (EMBO) Installation Grant (EMBO-2020-IG4734), and the Chan Zuckerberg Initiative Visual Proteomics Grant (vpi-0000000044). This research was supported by InFLAMES Flagship Programme of the Academy of Finland (decision number: 337531).

## AUTHOR CONTRIBUTIONS

Conceptualization, G.J.; Methodology, J.W.P, R.F.L., R.H., and G.J.; Formal Analysis, J.W.P., and G.J.; Production of the test datasets, S.G. and G.F.; Writing – Code: J.W.P, R.F.L., B.M.S. S., R.H., and G.J.; Writing – Original Draft, J.W.P and G.J.; Writing – Review and Editing, all authors; Visualization, J.W.P., and G.J.; Supervision, G.J.; Funding Acquisition, R.H. and G.J.

